# Dopamine-dependent cAMP dynamics in basal amygdala glutamatergic neurons

**DOI:** 10.1101/2021.09.03.458910

**Authors:** Andrew Lutas, Kayla Fernando, Stephen X. Zhang, Abhijeet Sambangi, Mark L. Andermann

**Affiliations:** Division of Endocrinology, Metabolism, and Diabetes, Beth Israel Deaconess Medical Center, Harvard Medical School, Boston, MA, USA, 02115

## Abstract

Dopaminergic inputs to basal amygdala (BA) instruct learning of motivational salience. Here, we investigated the dynamics of dopamine release and downstream signaling during multiple salient events occurring within tens of seconds. We established *in vitro* and *in vivo* real-time tracking and manipulation of cAMP – a key intracellular plasticity signal downstream of dopamine receptor activation. Optogenetically-evoked release of dopamine drove proportional increases in cAMP in almost all BA glutamatergic neurons, suggesting widespread actions of dopamine across neurons preferring positive or negative valence. These cAMP responses decayed more slowly than dopamine release, potentially extending the window of plasticity. cAMP levels accumulated following direct photostimulation of cAMP but not repeated stimulation of dopamine axons, due to potent depression of dopamine release. cAMP and protein kinase A (PKA) responses to repeated appetitive or aversive stimuli also exhibited pronounced depression. Thus, history-dependent dynamics of dopamine and cAMP may regulate learning of temporally clustered, salient stimuli.

## Introduction

The basolateral amygdala is critical for learning the valence of initially neutral sensory cues and guiding decisions to approach or avoid such cues (O’Neill et al., 2018; Zhang and Li, 2018). The neural plasticity that occurs during this associative learning involves dopamine, norepinephrine and other neuromodulators that bind to receptors on target cells to regulate cyclic adenosine monophosphate (cAMP) levels and synaptic plasticity (Bissière et al., 2003; Johansen et al., 2014; Tronson et al., 2006). During associative plasticity, the precise timing of dopamine-dependent cAMP signals is important (Handler et al., 2019; Steinberg et al., 2013; Yagishita et al., 2014). However, we understand much less about the dynamics of dopamine-evoked cAMP signals, which also depend on regulatory mechanisms that vary across states and across neuron types.

An increase in release of dopamine from VTA neurons projecting to the basal amygdala (BA) occurs during salient events of *both* positive and negative valence (Lutas et al., 2019). Despite receiving this common dopamine signal, individual BA glutamatergic neurons preferentially respond to *either* appetitive or aversive events (Lutas et al., 2019; O’Neill et al., 2018; Zhang and Li, 2018). These observations led us to hypothesize that dopamine release broadly facilitates plasticity by increasing cAMP levels in both appetitive- and aversive-preferring BA neurons during salient outcomes. Meanwhile, learning of the valence of the outcome associated with a cue may be achieved via other mechanisms involving calcium-dependent processes driven by distinct pathways relaying information about sensory cues and about positive and negative outcomes (Correia and Goosens, 2016).

We developed an approach to optogenetically control dopamine release from VTA axons in BA (VTA^DA→BA^) while simultaneously visualizing calcium or cAMP dynamics in BA glutamatergic neurons. To first test the sufficiency and timing-dependence of dopamine-evoked cAMP in instructing stimulus salience, we performed longitudinal imaging of calcium responses in BA neurons in awake mice across sessions in which one of two visual stimuli is paired with optogenetic stimulation of VTA^DA→BA^ axons. We found that optogenetic stimulation of dopamine axons was sufficient to drive the acquisition of stimulus responses in a subset of BA neurons across days, similar to the acquisition of responses to cues paired with natural appetitive and aversive outcomes (Lutas et al., 2019). We then investigated the immediate consequences of dopamine receptor activation on BA glutamatergic neurons by tracking intracellular cAMP production using a genetically-encoded fluorescent biosensor (Tewson et al., 2016). We found that exogenous dopamine and transient photostimulation of VTA^DA→BA^ axons elevated cAMP in most BA glutamatergic neurons. The proportion of BA neurons with dopamine-evoked increases in cAMP scaled with dopamine concentration, and all BA neurons showed elevated cAMP in the presence of high levels of exogenous dopamine. These findings suggest that while dopamine release does not determine the encoded valence of the conditioned stimulus, it may determine its salience by controlling the number of stimulus-responsive BA neurons.

We also found that transient dopamine release triggered cAMP increases lasting for over 30 seconds. This duration was primarily determined by cell-autonomous processes rather than by prolonged elevations in extracellular dopamine, since we could replicate the cAMP decay kinetics by circumventing dopamine receptors using a photoactivatable adenylate cyclase. Further, we found that presynaptic depression at VTA^DA→BA^ terminals limits the duration of elevated cAMP when dopamine release events are spaced tens (but not hundreds) of seconds apart. This depression in cAMP signaling was observed for repeated optogenetically-evoked dopamine release as well as for repeated delivery of unexpected palatable food or aversive tail shock. We also observed potent depression at the next stage in the signaling pathway, protein kinase A (PKA) activity, *in vivo* in response to repeated salient events. Thus, the learned salience associated with novel stimuli could be regulated by the magnitude of dopamine-evoked cAMP within the BA. Together, our results suggest that dopamine-evoked cAMP initiates a widespread permissive plasticity window that is correlated with stimulus salience and engages other intracellular signals in BA to control the acquisition of sensory responses to cues surrounding motivationally salient outcomes.

## Results

### Pairing visual cues with optogenetic stimulation of VTA^DA→BA^ axons results in cue-evoked calcium responses in BA neurons

Neurons in the basolateral amygdala acquire responses to previously meaningless visual cues when these cues are paired with salient appetitive or aversive outcomes (Lutas et al., 2019; O’Neill et al., 2018; Schoenbaum et al., 1999). To determine whether pairing dopamine release with an arbitrary visual stimulus was sufficient for BA neurons to acquire responses to this stimulus, we combined two-photon calcium imaging of the same individual BA neurons across several sessions with photostimulation of VTA^DA→BA^ axon terminals (Supplementary Figure 1A,B). We used two simple visual stimuli differing only in the orientation of a drifting grating. One of the stimuli (“Cue A”) was paired with photostimulation of VTA^DA→BA^ axon terminals, which occurred 200 ms after the visual stimulus offset, and the second (“Cue B”) was not paired with any outcome (Supplementary Figure 1C). An inter-trial interval (ITI) of 6 seconds was used to mirror our prior study that paired these visual stimuli with salient appetitive or aversive outcomes (Lutas et al., 2019). We first imaged BA neuron responses to the presentation of these visual stimuli prior to any manipulations. Only a small fraction of BA neurons was significantly responsive (1%; n = 14/1283 neurons from 7 mice) to the presentation of these arbitrary visual stimuli (Supplementary Figure 1D), consistent with prior work (Lutas et al., 2019). We then paired “Cue A” with brief photostimulation of VTA^DA→BA^ axon terminals across several daily sessions while tracking calcium responses in the same BA neurons. After three days of pairing, a larger fraction of BA neurons had become responsive to the paired visual stimulus (4%; n = 48/1199 neurons from 6 mice; Supplementary Figure 1D). Despite the randomized nature of stimulus presentations and the six-second ITI between trials, BA neurons that acquired responses to “Cue A” also acquired responses to the unpaired “Cue B” (Supplementary Figure 1E,F). As discussed below, such generalization may relate to sustained effects of transient VTA^DA→BA^ axon photostimulation. Nevertheless, responses to “Cue B” were weaker than those to “Cue A”, and neurons preferentially responded to “Cue A” rather than “Cue B” (Supplementary Figure 1G,H). Therefore, photostimulating VTA^DA→BA^ axons immediately following an arbitrary visual cue leads to the acquisition of cue-evoked responses in a subset of BA glutamatergic neurons.

### A cAMP sensor reveals spatially broad dopamine signaling across all BA glutamatergic neurons

To better understand the immediate effects of photostimulating VTA^DA→BA^ axons on postsynaptic neurons, we expressed a Cre-dependent, compact fluorescent biosensor for cAMP, cADDis (Tewson et al., 2016), in BA glutamatergic neurons via AAV injections into EMX1-Cre transgenic mice (Figure 1A,B). We tested the sensor functionality by applying the adenylate cyclase (AC) activator forskolin (Figure 1C) or dopamine (Figure 1D) to brain slices to stimulate intracellular cAMP production. Both forskolin and dopamine generated reliable decreases in mean fluorescence intensity, indicating increasing cAMP concentration (Tewson et al., 2016) (note that for visualization purposes, y-axes have been inverted so that a rise in cADDis signal reflects an increase in cAMP; Figure 1C).

**Figure 1.**
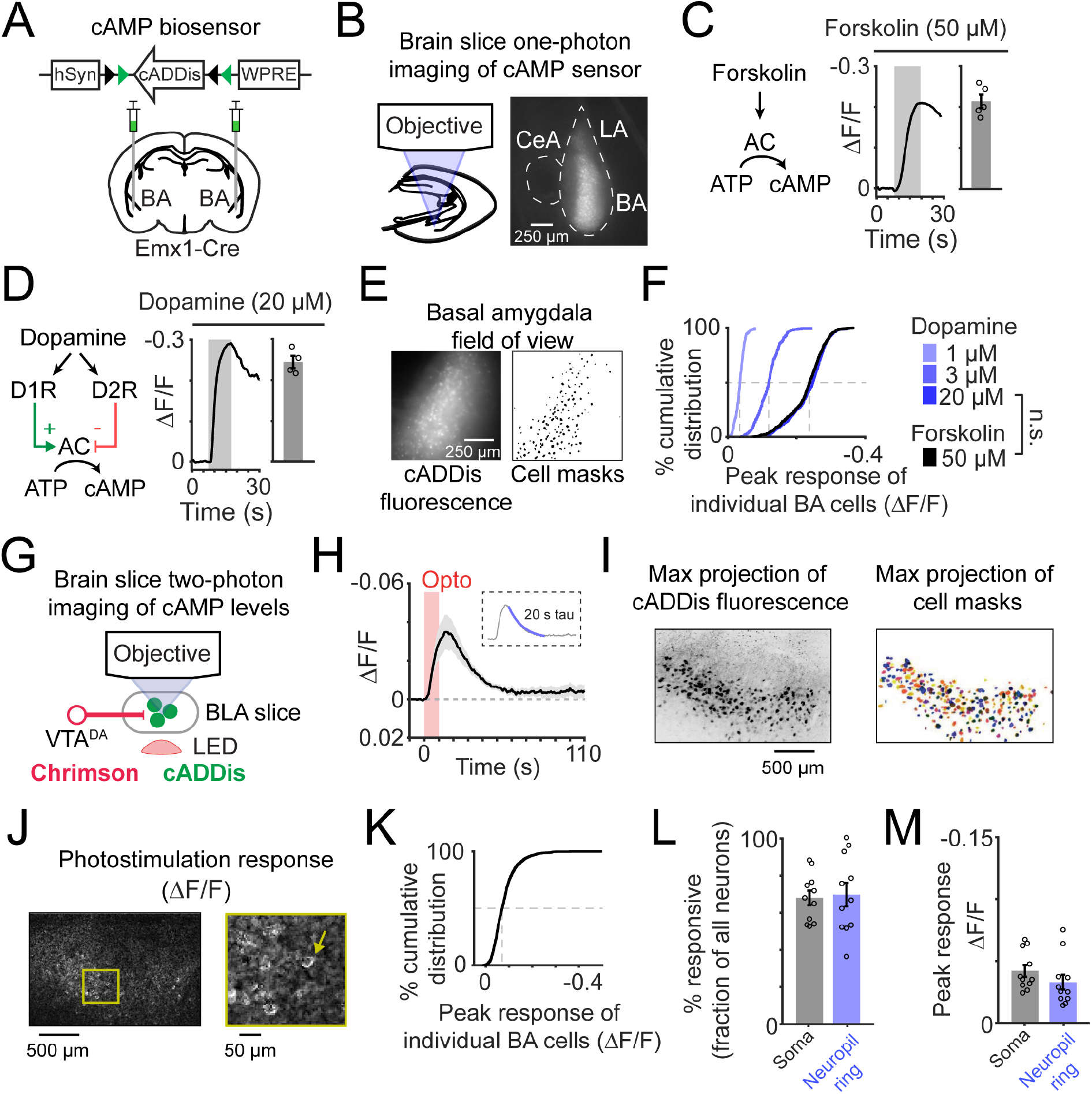
Endogenous dopamine release elevates cAMP in most basal amygdala glutamatergic neurons. **A)** *Top*: DNA construct for human Synapsin (hSyn) promotor driven, Cre-dependent expression of cADDis in neurons. Black and green triangles: Cre recombination sites. WPRE: woodchuck hepatis virus post-transcriptional regulatory element. *Bottom*: viral injection strategy for selective expression of cADDis in glutamatergic neurons of the basal amygdala (BA) using Emx1-Cre transgenic mice. **B)** *Left*: schematic of live cell epifluorescence imaging of cADDis in a brain slice. *Right*: representative image of amygdala brain slice expressing cADDis in BA neurons. CeA: central amygdala. LA: lateral amygdala. **C)** *Left*: forskolin-mediated activation of adenylate cyclase (AC) drives production of cAMP. *Middle*: example forskolin-evoked (50 μM) cADDis fluorescence change in BA region of a brain slice. Y-axis is inverted for clarity, as increases in cAMP decreased cADDis fluorescence. *Right*: peak response of individual slices. Mean ± s.e.m. n = 5 slices from 2 mice. **D)** *Left*: dopamine-mediated activation or suppression of adenylate cyclase (AC) via activation of either Type 1 (D1R) or Type 2 (D2R) dopamine receptors. *Middle*: example dopamine-evoked (20 μM) cADDis fluorescence change in BA. *Right*: peak response of individual slices. Mean ± s.e.m. n = 4 from 2 mice. **E)** *Left*: epifluorescence image of BA glutamatergic neurons expressing cADDis. *Right*: masks for individual cell somata. **F)** Average cumulative distribution of peak responses of individual BA neurons to dopamine and forskolin application. 1 μM dopamine: n = 440 neurons from 3 slices from 1 mouse. 3 μM dopamine: n = 294 neurons from 2 slices from 2 mice. 20 μM dopamine: n = 579 neurons from 4 slices from 2 mice. 50 μM forskolin: n = 597 neurons from 5 slices from 2 mice. ^n.s.^, p > 0.05. Kolmogorov-Smirnov test. **G)** Schematic of brain slice imaging of cADDis fluorescence and photostimulation-evoked dopamine release. VTA^DA→BA^ dopamine axons expressing Chrimson were stimulated using red-light illumination from below the slice. cAMP levels were determined by two-photon imaging of cADDis in BA glutamatergic neurons. **H)** Mean cADDis fluorescence in response to photostimulation of dopamine release (10 s duration; 15.5 Hz). Mean ± s.e.m. n = 12 slices from 4 mice. *Inset*: mean cADDis fluorescence with monoexponential fit overlaid (blue line). Fitted decay rate was 20 s. **I)** *Left*: maximum intensity projection of a volume (15 depths, ~10 μm apart) of cADDis fluorescence. The image color is inverted for clarity. *Right*: maximum projection of the color-coded cell masks of individual BA neurons from the imaged volume. **J)** Image of normalized difference in cADDis fluorescence between dopamine and baseline conditions in an example slice field of view. **K)** Cumulative distribution of peak cADDis responses of individual BA neurons following photostimulation of VTA^DA→BA^ axon terminals. n = 1697 neurons from 11 slices from 5 mice. **L)** Percentage of all neuronal somata (black) or neuropil rings surrounding somata (blue) per slice with significant cADDis responses following photostimulation of VTA^DA→BA^ axons. Mean ± s.e.m. n = 11 slices from 5 mice. **M)** Peak response of neuronal somata or neuropil rings with significant cADDis responses following photostimulation of VTA^DA→BA^ axons. Mean ± s.e.m. n = 11 slices from 5 mice.

We next used cADDis to estimate the fraction of BA glutamatergic neurons that *could* respond to dopamine. Since almost all BA glutamatergic neurons express Type 1 dopamine receptors (D1) (Lutas et al., 2019; Namburi et al., 2015; O’leary et al., 2020), while only few express Type 2 dopamine receptors (D2), dopamine should elevate cAMP in BA neurons. To directly determine the fraction of BA glutamatergic neurons that showed increases in cAMP in response to dopamine, we extracted fluorescence traces from the somatic regions of individual neurons (Figure 1E; see Methods). We first analyzed responses to application of forskolin as a measure of dopamine receptor-independent cAMP production. Forskolin resulted in cAMP increases in all BA glutamatergic neurons. We then found that application of a high concentration of dopamine (20 μM) potently elevated cAMP in *all* BA glutamatergic neurons, similar to forskolin, while lower concentrations drove weaker yet detectable increases in BA neurons (Figure 1F). This finding demonstrates a near-universal potential for dopamine signaling in BA glutamatergic neurons, and suggests that the diversity of valence preferences of nearby BA glutamatergic neurons (Lutas et al., 2019; O’Neill et al., 2018) may not to be due to differences in their ability to increase (or decrease) their cAMP levels in response to dopamine.

### Photostimulation of VTA^DA→BA^ axons transiently elevates cAMP in almost all BA glutamatergic neurons

While exogenous application of dopamine revealed a widespread increase in cAMP levels in BA glutamatergic neurons, physiological release of dopamine directly from axon terminals may be more spatially restricted. To evoke endogenous dopamine release, we targeted expression of a red-shifted channelrhodopsin (Chrimson) in VTA dopamine neurons using DAT-Cre transgenic mice (Bäckman et al., 2006; Klapoetke et al., 2014). To confirm that photostimulating VTA^DA→BA^ axons in brain slices reliably evoked dopamine release, we combined widefield photostimulation of Chrimson with two-photon imaging of a green fluorescent biosensor for dopamine, dLight1.1, that was expressed in BA glutamatergic neurons (Patriarchi et al., 2018) (Supplementary Figure 2A). Brief trains of optical stimulation (20 Hz; 5 s duration) generated transient elevations in dopamine that lasted for several seconds (Supplementary Figure 2B). Importantly, fluorescence changes in this D1-based dopamine sensor could be completely blocked by application of a D1 antagonist (SCH23390, 300 nM, Supplementary Figure 2C), confirming that the signals were mediated by dopamine release and not by artifacts related to optical stimulation. By comparing endogenous release of dopamine following optical stimulation of VTA^DA→BA^ axons to that following application of a near-saturating concentration of exogenous dopamine (cf. Figure 1), we found that the bulk endogenously released dopamine concentration was 5 times lower than near-saturating concentrations of dopamine (Supplementary Figure 2D), although the effective concentration at synaptic clefts may be higher.

We next tested whether we could visualize changes in cAMP levels in BA glutamatergic neurons in response to photostimulated dopamine release (Figure 1G). We crossed DAT-Cre and EMX1-Cre transgenic mice, which allowed for viral targeting of Chrimson to dopamine-releasing neurons in the VTA and cADDis to glutamatergic neurons in the BA. Photostimulation of VTA^DA→BA^ axon terminals combined with two-photon imaging of cADDis revealed transient increases in cAMP that peaked ~5 seconds later than dopamine, consistent with a delay between the rise in extracellular dopamine and the generation of intracellular cAMP in target neurons (Supplementary Figure 2E). The cAMP elevation remained above baseline for tens of seconds following optical stimulation of VTA^DA→BA^ axons – much longer than the duration of elevated extracellular dopamine (Figure 1H). The slower decay of cAMP was not due to prolonged activation of target dopamine receptors, as similar decay kinetics of evoked increases in cAMP were observed following direct photostimulation of intracellular cAMP using a novel blue-light activated adenylate cyclase (biPAC; Supplementary Figure 3A-C) (Zhang et al., 2021b). Thus, the release of dopamine onto BA neurons is transformed into a transient elevation in intracellular cAMP that extends in time by approximately five-fold compared to the elevation in extracellular dopamine (note that this is likely an underestimate of the duration of cAMP elevations given the low affinity of cADDis for cAMP, Kd ~ 10 μM). This extended elevation in cAMP across tens of seconds may reflect an extended window of plasticity that could result in generalization of plasticity to unpaired cues occurring within that time window (e.g. Supplementary Figure S1).

We next examined endogenous dopamine-evoked cAMP signals from individual BA neuron somata (Figure 1I,J; see Methods). Similar to cellular responses to exogenous dopamine application (Figure 1F), we detected widespread cAMP responses in over 70% of all BA glutamatergic neurons as well as in the surrounding neuropil largely composed of dendrites of these neurons (Figure 1K,L). While the magnitudes of the cAMP increases were small (Figure 1M), they were larger than those observed following 1 μM exogenous dopamine, a concentration that also drove cAMP increases across all BA neurons (Figure 1F). We observed similar magnitude responses in somatic and neuropil compartments, consistent with the finding that dopaminergic synaptic terminals contact dendrites of BA glutamatergic neurons (Muller et al., 2009) (Figure 1M). These findings indicate that most BA glutamatergic neurons express functional D1 receptors and can respond to axonal release of dopamine.

### Synaptic depression of dopamine release temporally restricts cAMP signals in BA neurons

We noticed that brief trains of optical stimulation (20 Hz; 2, 5, or 10 s duration) generated transient elevations in cAMP that increased sublinearly with pulse train duration (Figure 2A-C). This sublinearity may reflect presynaptic depression of dopamine, as has been observed for dopaminergic inputs to the striatum (Liu and Kaeser, 2019). Indeed, we observed that photostimulating VTA^DA→BA^ axons evoked dopamine release that scaled sublinearly with increasing duration of the 15 Hz pulse train (Figure 2D-F). We also observed a diminished magnitude of dopamine release when inter-trial intervals (ITIs) were 20 seconds long, but not when they were two minutes long (Figure 2G-I). We found that 20 second ITIs resulted in a 25% depression in dopamine release whereas 120 second ITIs did not result in depression (Figure 2H,I). This long-lasting depression of dopamine release is consistent with the depression of VTA^DA→BA^ axon-evoked glutamate co-released with dopamine in the BA (Lutas et al., 2019), and with evidence that VTA projections to the striatum also exhibit synaptic depression when ITIs are shorter than two minutes (Adrover et al., 2014).

**Figure 2.**
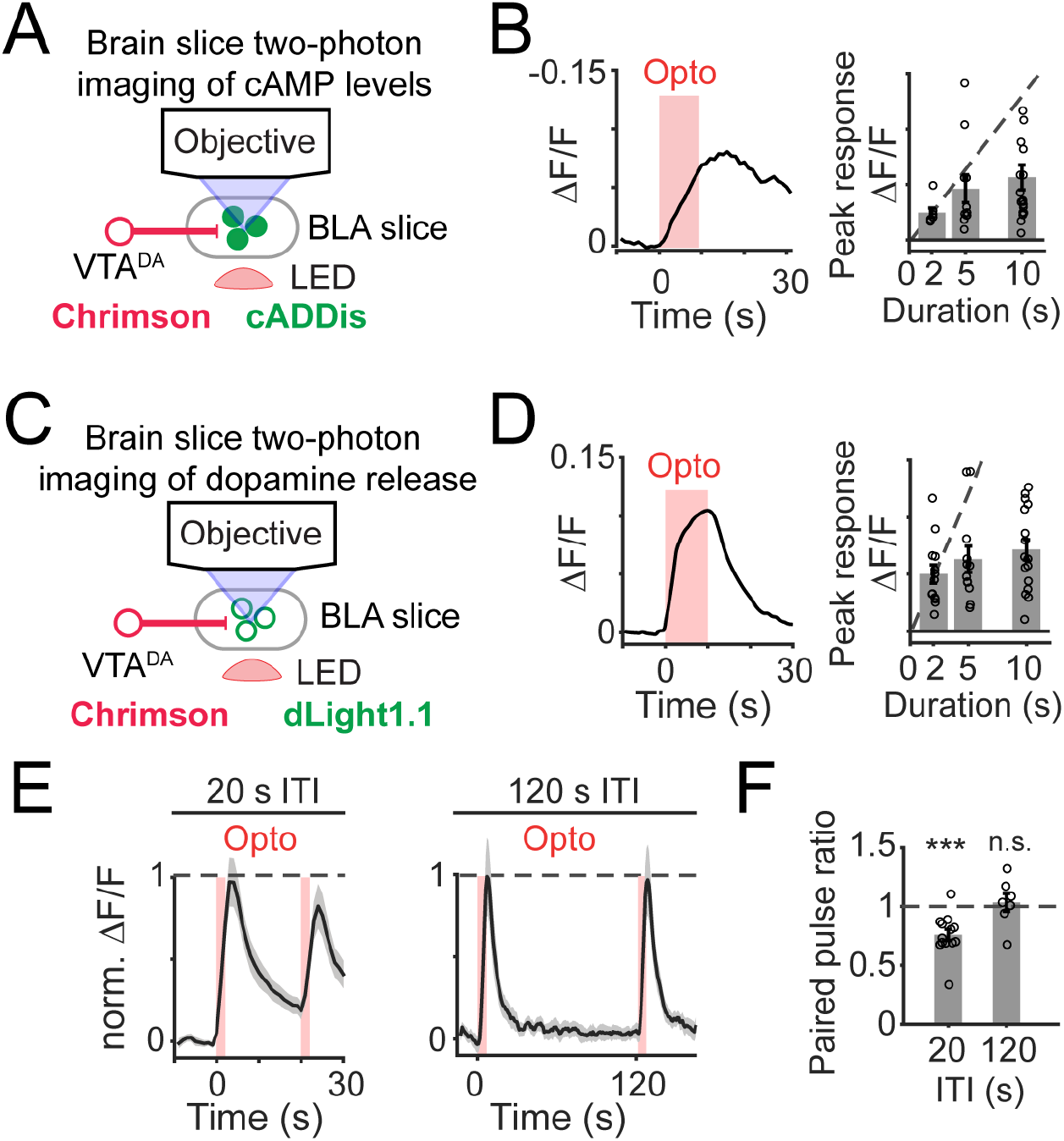
Synaptic depression of dopamine release restricts postsynaptic cAMP accumulation. **A)** Schematic of brain slice imaging of cADDis fluorescence and photostimulation-evoked dopamine release. VTA^DA→BA^ axons expressing Chrimson were stimulated using red-light illumination from below the slice. cAMP levels were determined by two-photon imaging of cADDis expressed in BA glutamatergic neurons. **B)** *Left*: example recording of cADDis fluorescence in response to a train of red-light pulses (10 s duration; 15.5 Hz). *Right*: peak response of individual brain slices to photostimulation of dopamine release for 2 s (n = 7 slices from 4 mice), 5 s (n = 11 slices from 5 mice), or 10 s (n = 13 slices from 5 mice). Mean ± s.e.m. **C)** Schematic of brain slice imaging of photostimulation-evoked dopamine release. Dopamine levels were determined by two-photon imaging of dLight1.1 expressed in BA. **D)** *Left*: example recording of dLight1.1 fluorescence in response to a train of red-light pulses (10 s duration; 15.5 Hz; 620 nm). *Right*: peak responses of average signal from individual brain slices to photostimulation of dopamine release for 2 s (n = 13 slices from 4 mice), 5 s (n = 12 slices from 4 mice), or 10 s (n = 16 slices from 4 mice). Mean ± s.e.m. **E)** *Left*: mean time course in response to a pair of photostimulation trials (2 s pulse train duration; 15.5 Hz; 20 s ITI) normalized to first peak. Mean ± s.e.m. n = 13 slices from 4 mice. *Right*: same as left but for 120 s ITI. n = 7 slices from 2 mice. **F)** Paired pulse ratio of peak response (second pulse train / first pulse train) for inter-trial intervals (ITIs) of 20 s (***, p = 0.0005, n = 13 slices from 4 mice) or 120 s (n.s., p = 0.57, n = 7 slices from 2 mice) duration. Two-sided Wilcoxon sign-rank. Mean ± s.e.m.

### *In vivo* cAMP dynamics in BA neurons following repeated photostimulation of VTA^DA→BA^ axons

We next asked whether cAMP dynamics in BA glutamatergic neurons exhibit similar characteristics *in vivo* as in our brain slice experiments. We used fiber photometry to track cADDis signals in BA glutamatergic neurons in response to photostimulation of Chrimson in VTA^DA→BA^ axon terminals (Figure 3A). We previously used this method of stimulation and recording to measure optical stimulation-evoked dopamine release of a similar magnitude as natural (shock-evoked) dopamine release (Lutas et al., 2019). We reliably detected cAMP production in BA neurons in response to optically induced dopamine release *in vivo* (Figure 3B). Similar to our slice measurements, we observed elevated cAMP levels lasting tens of seconds following brief optical stimulation of dopamine release (Figure 3B). Our protocol for *in vivo* photostimulation of VTA^DA→BA^ axons was well below saturating levels, as it drove weaker changes in cAMP than intraperitoneal injection of a high concentration of a dopamine Type 1 receptor agonist (SKF81297; 20 mg/kg; Supplementary Figure 4A). When we repeatedly photostimulated VTA^DA→BA^ axons every 30 s, we found that cAMP transients attenuated over time, such that within three minutes, they were profoundly reduced in magnitude (Figure 3C,D). This result led us to consider the possibility that the diminished responses were caused by presynaptic depression of dopamine release, as described in Figure 2.

**Figure 3.**
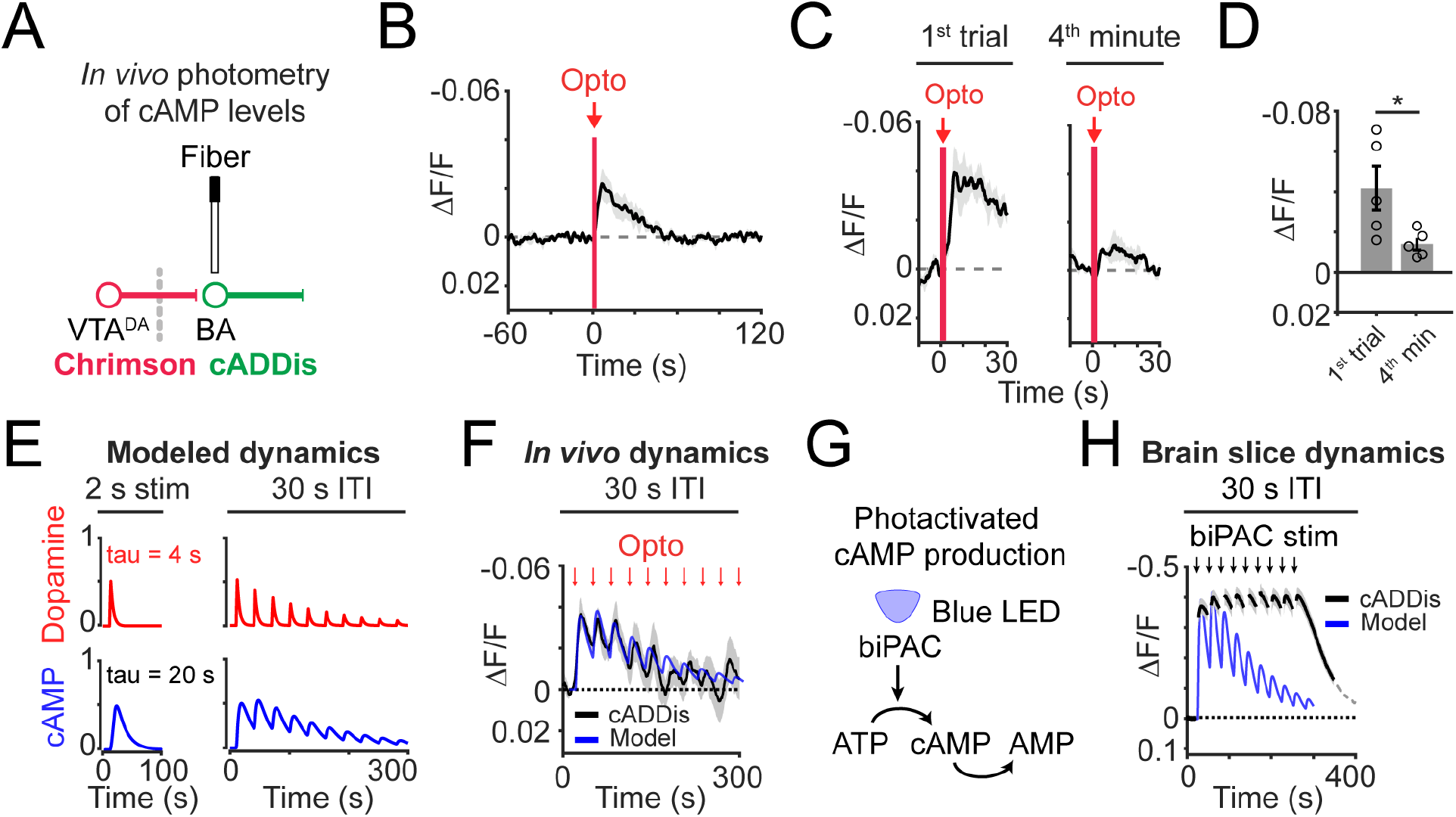
cAMP signals track photostimulated dopamine release *in vivo*. **A)** Schematic of *in vivo* photometry recordings of cADDis fluorescence and photostimulation of dopamine release. VTA^DA→BA^ dopamine axons expressing Chrimson were stimulated using red-light illumination via the same implanted fiber used to record cADDis signals from BA glutamatergic neurons. **B)** Mean time course of cADDis fluorescence in response to photostimulation of VTA^DA→BA^ axons (2 s duration; 20 Hz). n = 5 mice. Mean ± s.e.m. **C)** Mean time course of cADDis fluorescence of response to photostimulation of VTA^DA→BA^ axons (2 s duration; 20 Hz) for first trial of recording (*left*) or fourth minute of recording (*right:* average of 2 trials of 30 s duration). n = 5 mice. Mean ± s.e.m. **D)** Peak response following photostimulation on first trial *vs.* fourth minute (*, p = 03, n = 5 mice). Two-tailed paired t-test. Mean ± s.e.m. **E)** *Left*: simulation of dopamine (top; red) and cAMP (bottom; blue) dynamics for a single 2 s-duration activation. *Right*: simulation of stimulation of dopamine release at 30 s intervals. **F)** Mean time course of cADDis fluorescence (black line) of response to photostimulation of VTA^DA→BA^ axons (2 s duration; 20 Hz) every 30 s. Simulated cAMP dynamics (blue line) scaled to peak photometry amplitude. Mean ± s.e.m. n = 5 mice. **G)** Schematic of cAMP production using biPAC (blue-light stimulation of adenylate cyclase). **H)** Direct stimulation of cAMP production by blue-light activation of biPAC in brain slices (black; 2 s duration; 30 s ITI). Simulated cAMP dynamics (blue line) scaled to peak photometry amplitude. Mean ± s.e.m. n = 11 slices from 4 mice.

We explored this possibility by modeling cAMP dynamics in response to dopamine released at variable ITIs (30 s or 10 s; Figure 3E; Supplementary Figure 4B). We started with the assumption that the dynamics of intracellular cAMP in BA neurons following photostimulation of dopamine axons could be understood as a simple convolution of the dynamics of evoked dopamine concentration and of the decay in intracellular cAMP. We separately fit monoexponential decay functions to the post-peak response time courses from slice recordings for axon-evoked dopamine release and for cAMP evoked by direct, transient photostimulation of intracellular cAMP (dopamine: τ = 4 s; cAMP: τ = 20 s; see Methods). We then modeled the expected exponential attenuation in dopamine release with repeated stimulation based on our measurements of paired-pulse depression (Figure 2C; paired pulse ratio: 75%). Indeed, our simulation revealed that depression in dopamine release over trials is largely sufficient to result in a return of evoked cAMP levels to baseline within three minutes (Figure 3E; Supplementary Figure 4B).

We compared this simulation to *in vivo* fiber photometry of cADDis signals during the same protocol of photostimulation of VTA^DA→BA^ axons with 30- or 10-second ITIs. We found that endogenous cAMP dynamics mirrored those in our simulation (Figure 3F; Supplementary Figure 4C). To further assess whether a presynaptic rather than postsynaptic mechanism mediated the depression in evoked cAMP responses, we used biPAC to bypass the dopamine receptor and directly stimulate cAMP production (Figure 3G). When we photostimulated biPAC repeatedly every 30 seconds, cAMP levels remained persistently elevated through the duration of the recording (Figure 3H). Together, these results demonstrate that transient cAMP elevations such as those that likely occur during a rapid sequence of salient events do not accumulate across minutes, likely because of presynaptic depression of dopamine release.

### Repeated salient events evoke decreasing cAMP responses in BA neurons

Dopamine is naturally released in the BA during motivationally salient appetitive and aversive events (Lutas et al., 2019). We tested whether we could detect cAMP changes during unexpected delivery of appetitive palatable food in hungry mice or aversive unexpected tail shocks in sated mice (Figure 4A), both of which drive robust dopamine release in the BA (Lutas et al., 2019). On the first trial, we were able to detect significant changes in response to the consumption of unexpected food delivery or to the delivery of an unexpected tail shock (Figure 4B,C), albeit with a lower amplitude than that observed during photostimulation. Similar to our finding with repeated photostimulation of VTA^DA→BA^ axons, we found that repeated food delivery or tail shock resulted in attenuated cAMP transients, such that within 5 minutes, we were unable to detect significant evoked cAMP increases (Figure 4D,E). The depression of cAMP responses *in vivo* in response to tail shocks or food delivery that was repeated every 30 or 10 seconds was similar to expectations from our simulated dopamine-evoked cAMP signal (Figure 4F,G; Supplementary Figure 4D,E).

**Figure 4.**
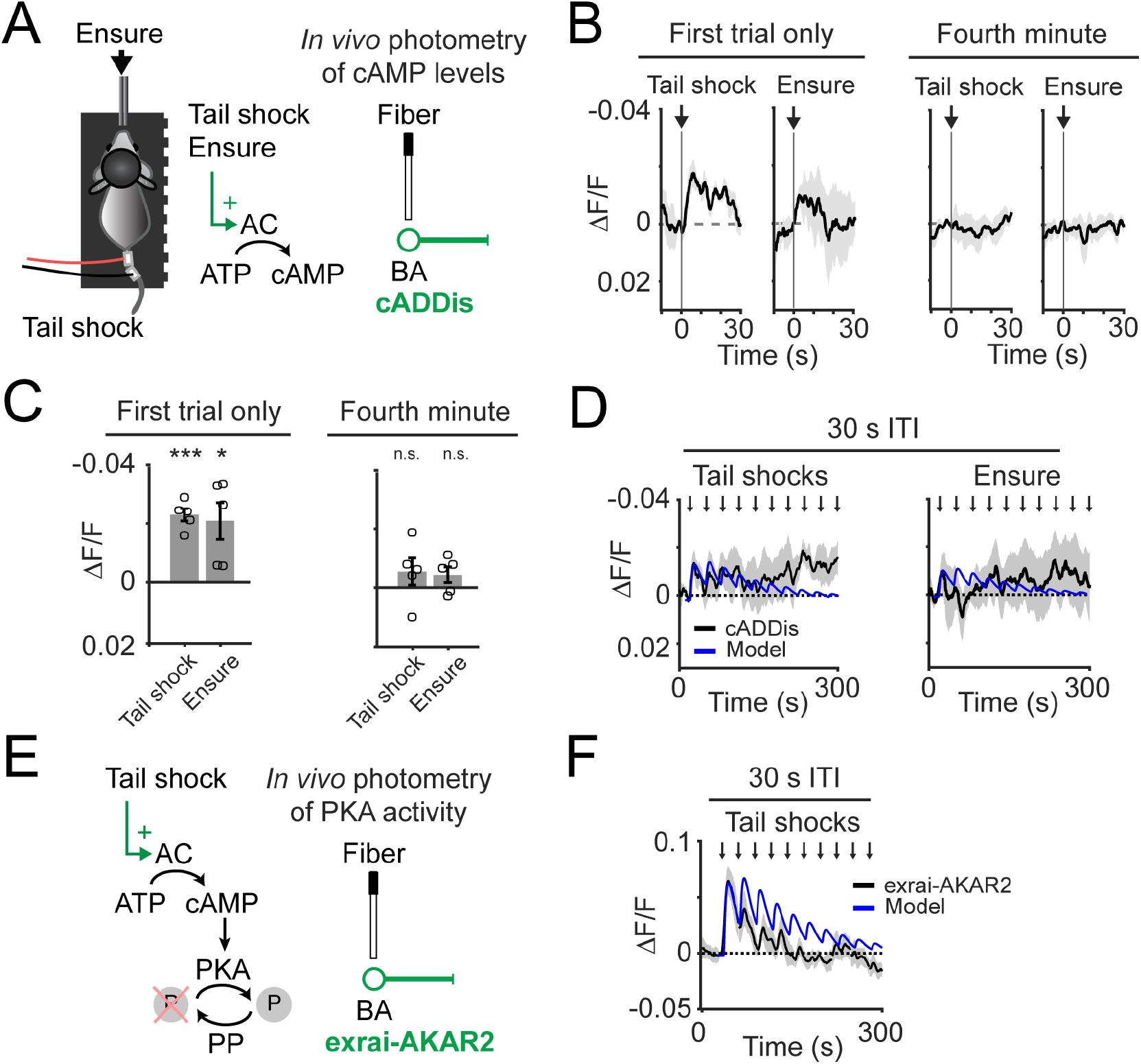
*In vivo* fiber photometry recordings of cAMP dynamics follow tail shock or Ensure delivery. **A)** *Left:* schematic of delivery of unexpected aversive tail shock or palatable food (Ensure) to an awake, head-fixed mouse. *Middle:* predicted consequences of tail shock or Ensure delivery on cAMP production in BA glutamatergic neurons. *Right:* schematic of *in vivo* photometry recordings of cADDis fluorescence. **B)** *Left*: mean time course of cADDis response to unexpected tail shock delivery (0.3 mA; 50 ms duration) or unexpected Ensure delivery via a solenoid (single 15 μL droplet) during first trial. *Right*: same as *left*, but for the fourth minute of recording. n = 5 mice. Mean ± s.e.m. **C)** Peak response following tail shock or Ensure delivery during the first trial (*left*) or fourth minute (*right*). ***, p = 0.0004; *, p = 0.03, ^n.s.^, p > 0.05, n = 5 mice. Two-tailed t-test. Mean ± s.e.m. **D)** *Left:* mean time course of cADDis fluorescence in response to unexpected tail shock delivery (0.3 mA; 50 ms duration) every 30 s. *Right:* mean time course of cADDis fluorescence in response to unexpected Ensure delivery (single 15 μL droplet) every 30 s. Recording normalized to baseline period before first trial. Mean ± s.e.m. n = 5 mice. Simulated cAMP dynamics (blue line; based on model shown in Figure 3E) scaled to peak photometry amplitude. **E)** *Left:* predicted consequences of tail shock on protein kinase A (PKA) activity. Dephosphorylation by protein phosphatases (PP) also indicated. *Right:* schematic of *in vivo* photometry recordings of fluorescence intensity of exrai-AKAR2 (biosensor of PKA activity, as measured by phosphorylation of PKA substrate peptide). **F)** Mean time course of exrai-AKAR2 fluorescence in response to unexpected tail shock delivery (0.3 mA; 50 ms duration) every 30 s. Recording normalized to baseline period before first trial. Mean ± s.e.m. n = 6 mice. Simulated cAMP dynamics (blue line; based on model shown in Figure 3E) scaled to peak photometry amplitude.

Finally, we addressed whether depression of cAMP signals also resulted in depression rather than accumulation of protein kinase A (PKA) activity - a key downstream regulator of synaptic plasticity (Figure 4H). We use fiber photometry and a fluorescent sensor of PKA activity, exrai-AKAR2 (Zhang et al., 2021a), to detect *in vivo* responses to tail shocks (Figure 4H). We found that unexpected tail shock delivery elicited responses in the exrai-AKAR2 photometry signals that were more pronounced than in our cADDis recordings (Figure 4I). We then tested if repeated delivery of tail shocks every 30 s produced a depressing exrai-AKAR2 response. Indeed, consistent with the depression in the evoked cAMP response magnitude, we found that exrai-AKAR2 responses depressed with repeated unexpected tail shocks (Figure 4I). While exrai-AKAR2 reports the ratio of phosphorylation by PKA and dephosphorylation by protein phosphatases, our earlier findings of depressing cAMP responses to repeated dopamine axon photostimulation suggests a weakening of cAMP-mediated PKA activation. However, we cannot fully rule out the possibility that increases in protein phosphatase activity also contribute to the depression in exrai-AKAR2 signal. Together, our characterization of *ex vivo* and *in vivo* cAMP and PKA sensor responses following endogenous dopamine release reveals temporal constraints on sustained cAMP elevation and PKA phosphorylation of targets in BA neurons. These temporal constraints likely affect the potency of associative plasticity of novel vs. rapidly repeated salient events.

## Discussion

The release of neuromodulators including dopamine in basal amygdala is critical for cue-outcome association learning (Johansen et al., 2014; Tang et al., 2020). Dopamine receptor activation, via cAMP dynamics, may strengthen synaptic inputs to enhance the salience of stimuli paired with either appetitive or aversive outcomes (Lutas et al., 2019). Here, we establish an imaging approach to investigate this process in BA glutamatergic neurons using optical tools to monitor cAMP dynamics and to evoke dopamine release or drive direct cAMP production. We demonstrate that exogenous dopamine as well as dopamine release from VTA^DA→BA^ axon terminals can increase cAMP in all BA glutamatergic neurons. Given that BA glutamatergic neurons segregate into appetitive- or aversive-preferring neurons, this widespread dopamine-evoked cAMP signal ignores the boundaries imposed by valence-specific teaching signals.

We found that during stimulation protocols that mimic temporally clustered, salient events, VTA^DA→BA^ axons initially drive cAMP in BA neurons *in vivo*. However, synaptic depression of these VTA^DA→BA^ axons limits the temporal window of dopamine-related accumulation of cAMP. In contrast, direct, repeated photostimulation of cAMP production that bypasses the dopamine receptor does not show this depression, and instead drives a persistent elevation in cAMP levels. These and other findings confirm that depression of dopaminergic input and actions of phosphodiesterases (Supplementary Figure 3C) control the accumulation of cAMP in BA neurons, thereby potentially regulating windows of plasticity in this region. More generally, our findings regarding the dynamics of dopamine release and cAMP accumulation in BA *in vitro* and *in vivo* provide a platform for linking the dynamics of intracellular biochemical signals with the dynamics of within-trial, across-trial, and across-day plasticity and learning.

### Synaptic depression of dopamine release in BA restricts the window of elevated cAMP when release events are clustered in time

We found a potent and long-lasting depression of dopamine-evoked cAMP transients. We showed that synaptic depression of dopamine release occurs at VTA^DA→BA^ terminals and that modeling this synaptic depression fully captures the weakened cAMP signals we recorded *in vivo* in response to photostimulated dopamine release. While to our knowledge this property had not been measured previously at the VTA^DA→BA^ synapse, similar characteristics have been observed in dopamine neuron projections to the dorsal and ventral striatum (Adrover et al., 2014). Synaptic depression at these mesostriatal dopaminergic synapses is mediated by several factors including the activation of presynaptic Type 2 dopamine receptors following release of dopamine, and by depletion of the synaptic vesicle pool (Liu and Kaeser, 2019). We suspect that similar mechanisms mediate the minutes-long depression we observed in VTA^DA→BA^ projections, both in slices and *in vivo*. We demonstrated that this depression greatly impairs the ability of dopamine axon activation to maintain elevated postsynaptic cAMP levels. Thus, depression of dopamine release may impose temporal constraints on synaptic plasticity when salient events are clustered in time. It will be interesting to test whether clustering together or spacing apart salient events using the temporal information obtained from cAMP measurements influences learning rates or memory consolidation in amygdala-dependent tasks, as has been observed in *Drosophila* (Jacob and Waddell, 2020). Moreover, as certain medications (e.g. methylphenidate) as well as drugs of abuse (e.g. cocaine) can enhance dopamine synaptic depression (Adrover et al., 2014), future studies can investigate how these drugs influence the temporal properties of amygdala plasticity.

### Is a permissive plasticity signal from VTA^DA→BA^ axons broadcast widely or only to a subset of BA neurons?

A major goal of our efforts to image cAMP in BA neurons was to understand whether dopamine, which is released from individual VTA^DA→BA^ axons following both appetitive and aversive outcomes (Lutas et al., 2019), drives increases in cAMP in all BA glutamatergic neurons. This could allow for dopamine to encode the motivational salience of an event by scaling the proportion of BA neurons that undergo a rise in cAMP levels. If different BA neurons exhibit varying sensitivities to dopamine, greater dopamine release during events with higher motivational salience should result in a greater percentage of BA neurons with dopamine-evoked changes in downstream signaling. We found that exogenous dopamine could elevate cAMP in *all* BA glutamatergic neurons, and photostimulation of endogenous release from dopamine axons drove detectable cAMP responses in most BA glutamatergic neurons in slices. On the one hand, this is likely to be a conservative estimate of affected neurons, given the limited sensitivity of our cAMP sensor (see below). On the other hand, we found that endogenous release of dopamine in response to palatable food and unexpected tail shocks were weaker than the photostimulated release, consistent with the interpretation that fewer BA neurons may have elevated cAMP following natural release of dopamine. The depression in dopamine-evoked cAMP we observed could further limit the magnitude of cAMP in some BA neurons. Thus, if different BA neurons have varying magnitudes of dopamine-evoked cAMP, the size of the BA neuronal population that exhibits plasticity could be limited to those neurons with the largest cAMP response. These results suggest that dopamine may determine the percentage of BA neurons that are plastic at a given time, thereby scaling learning rates with the motivational salience of expected appetitive and aversive outcomes.

### Potential limitations imposed by the sensitivity of the cAMP sensor

The affinity of the cAMP sensor we used is in the low micromolar range (Tewson et al., 2016), which means that changes in cAMP concentration in the nanomolar range would be outside of this sensitivity range. Especially in the case of our *in vivo* fiber photometry measurements, we may have missed changes in cAMP in response to unexpected delivery of palatable food or tail shocks. We overcame this limitation via additional experiments using a fluorescent biosensor of PKA activity (Zhang et al., 2021a), which had improved signal-to-noise in response to unexpected tail shocks and revealed clear depression with repeated salient events. An additional limitation is that our sensor measurements were not targeted specifically to dendritic compartments where cAMP increases may be concentrated, and thus our bulk measurements across somatic and dendritic compartments may have limited our sensitivity further. Nevertheless, presynaptic depression of dopamine release likely limits cAMP and PKA signals both in dendritic and somatic compartments. Future studies using higher affinity cAMP sensors (Klarenbeek et al., 2015) as well as sensors targeted to dendritic spines (e.g. by fusing with PSD-95) should allow for further real-time interrogation of these signals in response to endogenous salient events.

### Conclusions and future directions

We have established an imaging platform using reporters and actuators of intracellular cAMP that allowed for direct interrogation of dopamine-dependent cAMP signals in BA glutamatergic neurons *in vitro* and *in vivo*. Future studies can employ this approach to investigate cAMP dynamics in BA neurons in pre-clinical settings, such as following acute and chronic exposure to addictive substances, or to stressful, traumatic events. In addition, many other critical neuromodulatory signals in the BA (e.g., norepinephrine, PACAP, and serotonin) act via influences on cAMP. A similar imaging approach could be used to investigate how these additional neuromodulators affect cAMP levels alone and in conjunction with dopamine signaling. Another key future direction will involve examination of cAMP dynamics in non-glutamatergic targets of VTA dopamine neurons in BA, such as parvalbumin-positive interneurons (Chu et al., 2012; Pinard et al., 2008). In conclusion, we have revealed important temporal and spatial characteristics of dopamine actions via cAMP signaling in BA, which have informed hypotheses about the role of dopamine during associative learning. Importantly, while most *in vivo* studies of cellular plasticity have focused on changes in electrical and calcium activity, continued optimization of tools to detect molecular signaling cascades will expand our understanding of the underlying biochemical signals that control neural circuit plasticity.

## Acknowledgements

We thank members of the Andermann lab including K. Evans, R. Essner, N. Nguyen, K. McGuire, Dr. H. Kucukdereli, Dr. O. Amsalem, Dr. J.S. Alvarado for helpful feedback on the manuscript. We thank V. Flores-Maldonado for assistance with mouse colony care. We thank Dr. J. Madara for assistance with brain slice preparation. Boston Children’s Hospital Viral Core provided viral packaging services. We thank Drs. Ingie Hong and Richard Huganir for sharing exrai-AKAR2 virus. Authors were supported by an NIH F32 DK112589, a Davis Family Foundation Award, and a Boston Nutrition Obesity Research Center Pilot grant (A.L.), a Lefler Fellowship (S.X.Z.), NIH R01 DK109930, DP1 AT010971, DP1 AT010971-02S1, R01 MH12343, the McKnight Foundation, the Klarman Family Foundation, and the Harvard Brain Science Initiative Bipolar Disorder Seed Grant, supported by Kent and Liz Dauten (M.L.A).

The authors declare no conflicts of interest.

## Author contributions

A.L. and M.L.A. conceived the project and wrote the manuscript. A.L. designed and performed brain slice and two-photon imaging. A.L., K.F. and A.S. conducted fiber photometry recordings. A.L., K.F., A.S. and S.X.Z. performed surgical procedures. A.L. and S.X.Z. designed experiments for photostimulation concurrent with two-photon imaging of acute brain slices. A.L. analyzed all data.

---

**Supplementary Figure 1.**
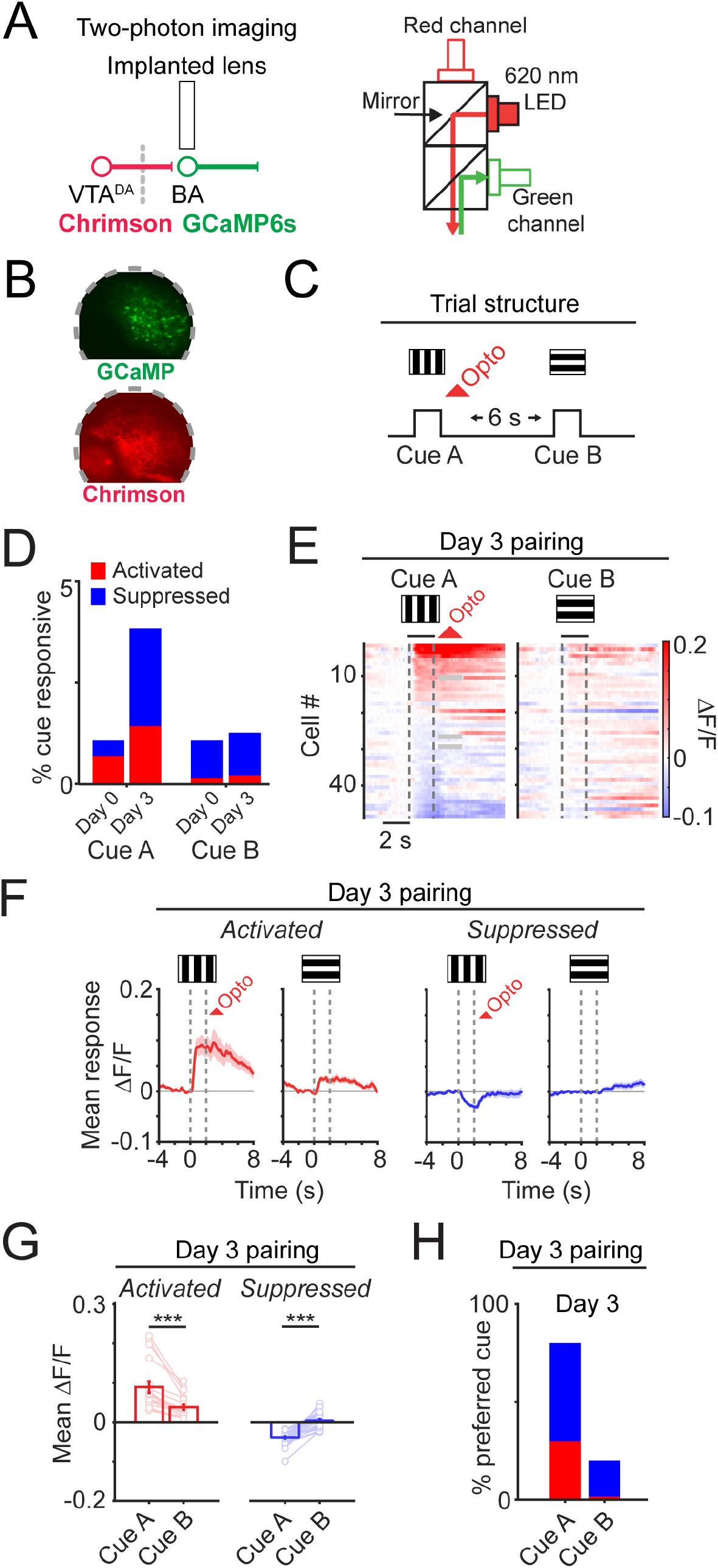
Two-photon calcium imaging during association of visual stimulus and photostimulation of VTA^DA→BA^ axons. **A)** *Left*: schematic of *in vivo* two-photon imaging of GCaMP6s fluorescence and photostimulation of dopamine release. VTA^DA→BA^ dopamine axons expressing Chrimson were stimulated using red-light illumination via the same implanted lens used to record GCaMP6s signals from BA glutamatergic neurons. *Right*: schematic of two-photon microscope optical path for red-light illumination and concurrent collection of green GCaMP6s fluorescence emission. **B)** *Top*: example image of GCaMP6s expression in BA glutamatergic neurons. *Bottom*: image of VTA^DA→BA^ axons expressing Chrimson-tdTomato in the same field of view. **C)** Diagram of trial structure design for pairing one visual stimulus (vertically oriented drifting bars, “Cue A”; 2 s duration) followed 200 ms later by photostimulation (5 mW, 2 s duration; 15.5 Hz). A second visual stimulus (horizontally oriented drifting bars, “Cue B”; 2 s duration) was not directly paired with photostimulation. **D)** Percentage of all neurons with significant cue responses on “Day 0” before photostimulation (Cue A: 14/1283 neurons; Cue B: 14/1283 neurons from 7 mice) and following three days of photostimulation (Cue A: 48/1199 neurons; Cue B: 12/1199 neurons from 6 mice). **E)** Heatmap with rows depicting mean response of BA neurons (n = 48 neurons, 14 fields of view from 6 mice) during presentation of each cue on the third day of photostimulation. **F)** Mean time course of responses to visual stimuli and photostimulation across all BA neurons that were significantly activated (red; n = 18 neurons) or suppressed (blue; n = 30 neurons) by “Cue A”. **G)** Mean response to “Cue A” or “Cue B” for activated (*left*, n = 18 neurons from 6 mice, ***, p < 0.001) and suppressed neurons (*right*, n = 30 neurons from 6 mice, ***, p < 0.001). Error bars: s.e.m. across neurons. Two-sided Wilcoxon sign-rank. **H)** Percentage of cue-responsive neurons preferring (i.e., maximally responsive to) a given cue following three days of photostimulation (Cue A: 48/60 neurons; Cue B: 12/60 neurons).

**Supplementary Figure 2.**
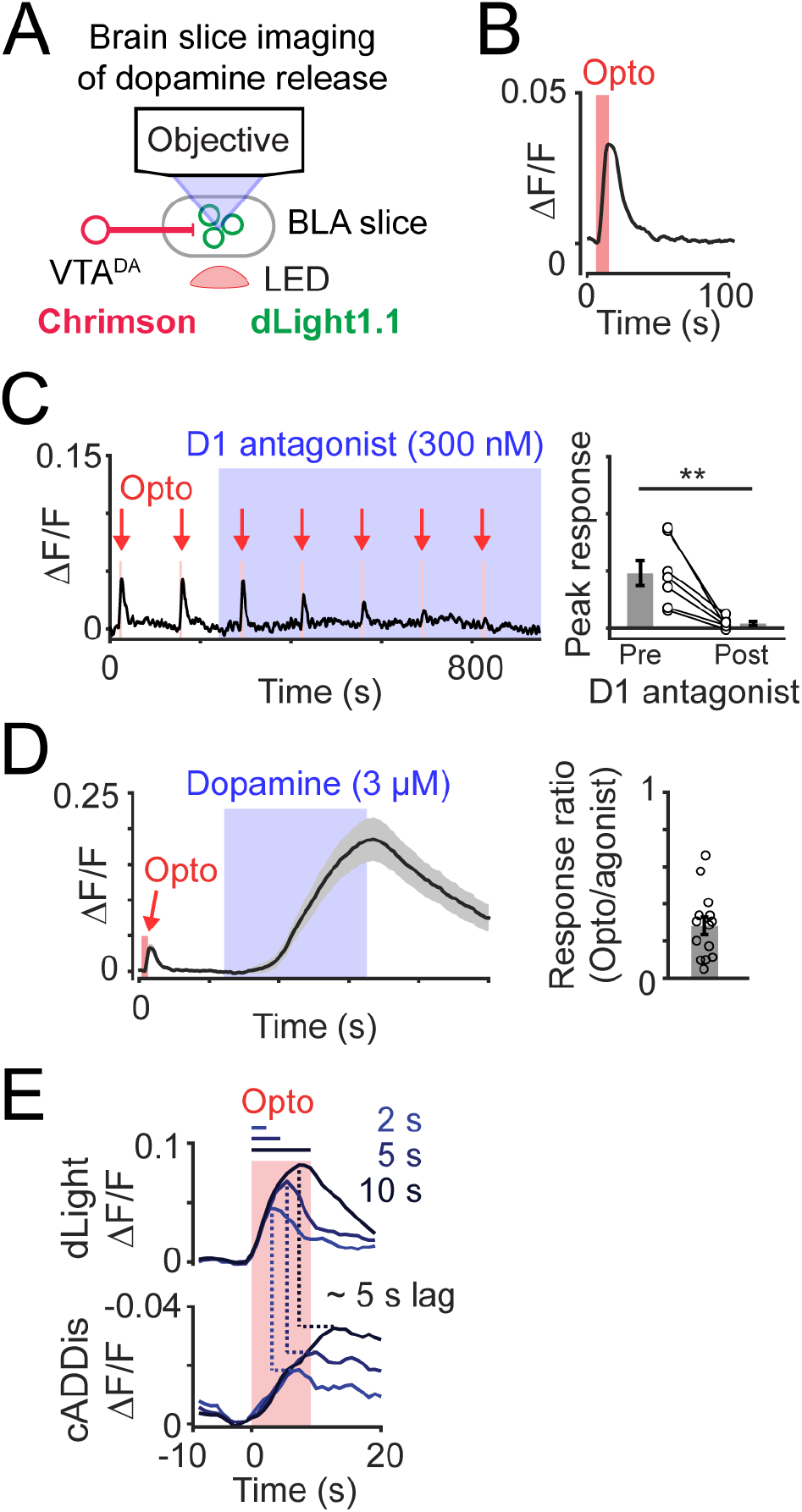
Photostimulation of VTA^DA→BA^ axons transiently elevates dopamine in BA. **A)** Schematic of brain slice imaging of photostimulation-evoked dopamine release. VTA^DA→BA^ dopamine axons expressing Chrimson were stimulated using red-light illumination from below the slice. Dopamine levels were determined by two-photon imaging of dLight1.1 expressed in BLA neurons. **B)** Mean dLight1.1 fluorescence during photostimulation of dopamine release (5 s duration; 15.5 Hz). n = 14 slices from 5 mice. Mean ± s.e.m. **C)** Application of dopamine D1 receptor antagonist (SCH23390; 300 nM) blocks photostimulation-evoked dLight1.1 signal. *Right*: peak photostimulation-evoked dLight1.1 response before and after antagonist. **, p = 0.006, n = 7 slices from 2 mice. Two-tailed paired t-test. Mean ± s.e.m. **D)** *Left*: mean time course in response to photostimulation (5 s duration; 15.5 Hz) followed by application of dopamine (3 μM). Mean ± s.e.m. *Right*: response ratio of exogenous dopamine vs. photostimulation-evoked dopamine release. n = 14 slices from 5 mice. Mean ± s.e.m. **E)** *Top*: mean time course of dLight1.1 fluorescence change in response to photostimulation of VTA^DA→BA^ axons for 2, 5, or 10 s. *Bottom*: mean time course of cADDis fluorescence changes in response to photostimulation of VTA^DA→BA^ axons for 2, 5, or 10 s.

**Supplementary Figure 3.**
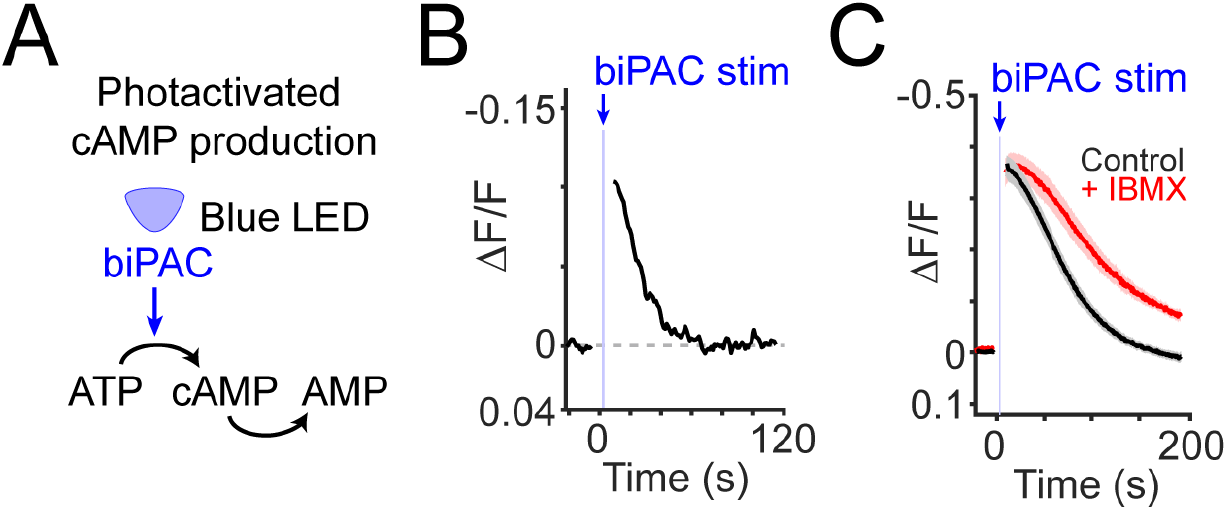
Photostimulation of direct cAMP production drives persistently elevated cAMP. **A)** Schematic of cAMP production using biPAC (blue-light stimulation of adenylate cyclase). **B)** Example recording of cADDis fluorescence during biPAC photostimulation (2 s duration; continuous illumination). Recording performed at 32°C. **C)** Stimulation of biPAC (2 s duration) in control slices (black; n = 13 slices from 4 mice) or in the presence of a phosphodiesterase inhibitor (IBMX, 100 μM; red; n = 4 slices from 4 mice). Recording performed at room temperature. Mean ± s.e.m.

**Supplementary Figure 4.**
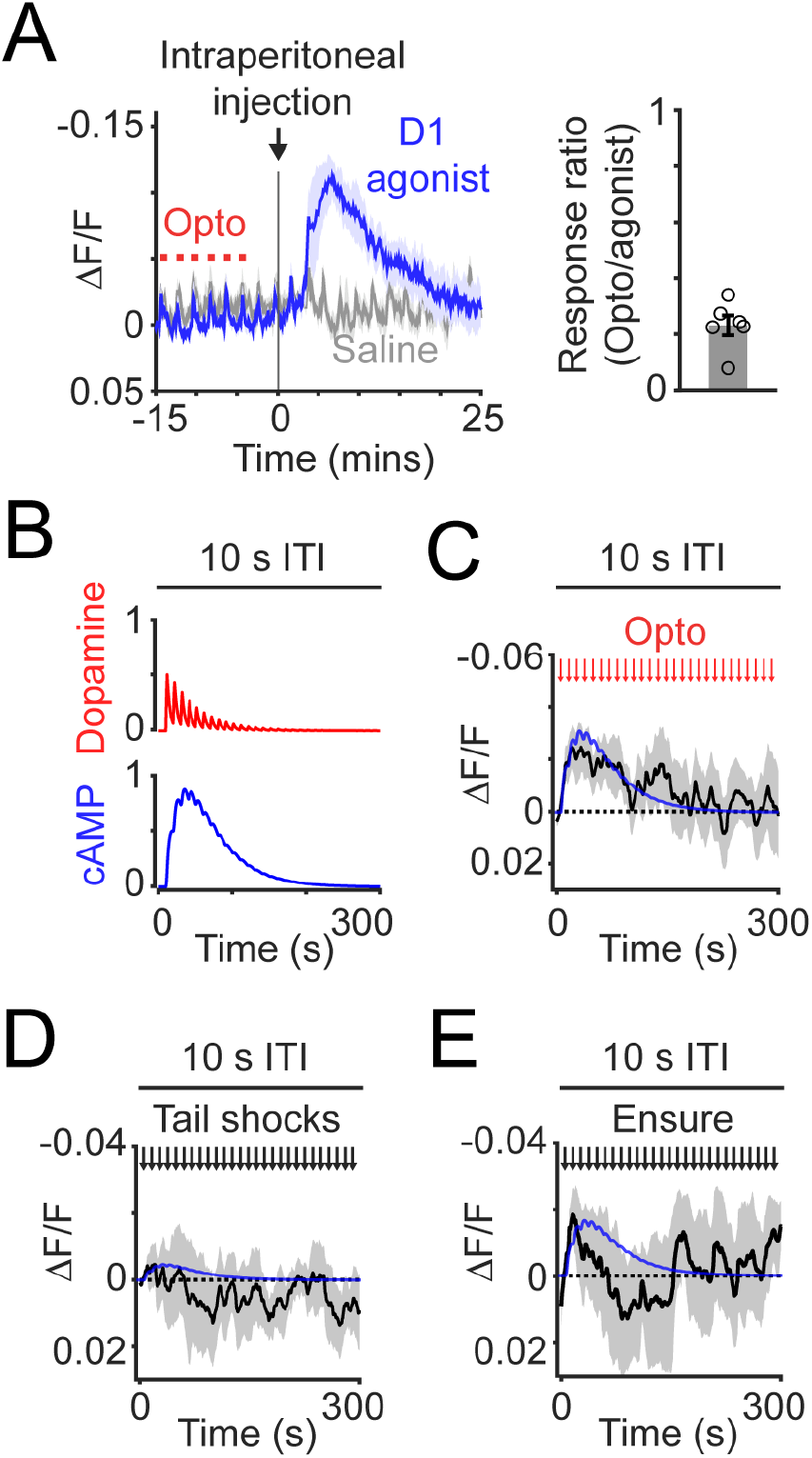
Additional fiber photometry of cADDis. **A)** *Left*: mean time course of cADDis fluorescence in response to intraperitoneal injection of a Type 1 dopamine receptor (D1) agonist (blue; SKF81297, 20 mg/kg, 150 μL; n = 6 mice) or saline (gray, 150 μL; n = 5 mice). Right: response ratio of injected D1 agonist to photostimulated dopamine response. n = 6 mice. Mean ± s.e.m. **B)** Simulation of repeated 2 s-duration stimulations of dopamine release occurring with 10 s inter-trial intervals (ITI). **C)** Mean time course of cADDis fluorescence (black line) of response to photostimulation of VTA^DA→BA^ axons (2 s duration; 20 Hz) every 10 s. Simulated cAMP dynamics (blue line) scaled to peak photometry amplitude. Mean ± s.e.m. n = 5 mice. **D)** Mean time course of cADDis fluorescence in response to unexpected tail shock delivery (0.3 mA; 50 ms duration) delivered every 10 s. Mean ± s.e.m. n = 4 mice. Simulated cAMP dynamics (blue line) scaled to peak photometry amplitude. **E)** Mean time course of cADDis fluorescence in response to unexpected Ensure delivery (single 15 μL droplet) delivered every 10 s. Simulated cAMP dynamics (blue line) scaled to peak photometry amplitude. Mean ± s.e.m. n = 3 mice.

## Lead contact

Further information and requests for resources and reagents should be directed to and will be fulfilled by the lead contact, Mark Andermann (manderma [at] bidmc.harvard.edu).

## Materials availability

For inquiries regarding cADDis plasmid and virus generated during this study, please contact Montana Molecular.

mKate2-biPAC plasmid DNA generated during this study and another study (Zhang et al., 2021) will be deposited to Addgene [catalog # 169127].

## Code and data availability

Analysis scripts were written in Matlab and used built-in functions and previously published code. No novel code was generated for the purpose of this study. Data is available upon request.

## EXPERIMENTAL MODEL AND SUBJECT DETAILS

### Animals

All animal care and experimental procedures were approved by the Institutional Animal Care and Use Committee at Beth Israel Deaconess Medical Center. Animals were housed in a 12-hour-light/12-hour-darkness environment with standard mouse chow and water provided *ad libitum*, unless specified otherwise. For both *in vitro* and *in vivo* experiments, adult male and female mice between the ages of 9 to 20 weeks were used for this study.

## METHOD DETAILS

### Stereotaxic surgeries

Viral injections, fiber implantations, and GRIN lens implantations were performed as described in (Lutas et al., 2019) with the following specifications and modifications.

For targeting expression into BA glutamatergic neurons, mice (8 – 12 weeks of age) expressing Cre recombinase driven by the Emx1 promoter (EMX1-Cre) were injected at (relative to Bregma): anteroposterior (AP): − 1.6 mm; dorsoventral (DV): − 4.8 mm; mediolateral (ML): ± 3.3 mm. For targeting expression into VTA dopamine neurons, DAT-Cre mice were injected at AP: − 3.2 mm; DV: − 4.5 mm; ML: ± 0.4 mm.

For *in vivo* experiments using the cAMP sensor cADDis, we unilaterally injected AAV1-hSyn-DIO-GreenDownward-cADDis (Children’s Hospital Vector Core; 300 nL). For cADDis or dLight1.1 (AAV1-hSyn-dLight1.1; Children’s Hospital Vector Core; 300 nL) brain slice imaging experiments, injections were targeted bilaterally to increase the number of useable slices. For *in vivo* calcium imaging experiments, we unilaterally injected AAV1-hSyn-FLEX-GCaMP6s (Addgene; 300 nL).

For fiber photometry experiments, optic fibers (400 μm diameter core; multimode; NA 0.48; 5.0 mm length; Doric Fibers) were implanted over BA at (relative to Bregma): AP: −1.6 mm; DV: −4.7 mm; ML: 3.3 mm.

For *in vivo* BA cell body imaging of GCaMP6s, mice were implanted with a singlet gradient index (GRIN) lens (GRINtech, NEM-100-25-10-860-S-1.0p; 1.0 mm diameter; 9 mm length; 250 μm focal distance on brain side at 860 nm, (NA 0.5); 100 μm focal distance on air side (NA 0.5); non-coated). GRIN lens implantation coordinates for cell body imaging of BA neurons in EMX1-Cre transgenic mice (relative to Bregma): AP: −1.6 mm, ML: 3.2 mm, DV: −4.8 mm. In order to ensure a snug fit for the lens to reduce brain motion and to increase surgical survival rate, we pre-set the insertion tracks by lowering a syringe needle with a slightly narrower diameter (20-gauge, 0.9 mm) to a depth of 0.1 mm above the final depth of the lens.

### Widefield epifluorescence imaging of acute brain slices

Following 3 – 5 weeks of expression, acute slices were prepared as described in Lutas *et al.,* 2019, and widefield fluorescence imaging was performed on an upright microscope (Axioskop 2 plus; Zeiss) equipped with an sCMOS II camera (Prime, Photometrics). Fluorescence excitation for imaging was achieved using a 470 nm LED (Thorlabs). Image acquisition was performed using ImageJ Micro-manager (Edelstein et al., 2014). Image acquisition frame rate was 2 Hz for cADDis fluorescence imaging. A 10x (Olympus) or 20x (Zeiss) objective was used for all imaging experiments. During imaging, slices were continuously superfused (flow rate: 2-5 ml/min) with oxygenated (95% O_2_ and 5% CO_2_) artificial cerebrospinal fluid (ACSF) at room temperature. To prevent oxidation of dopamine, 50 μM Na-metabisulfite was included in all ACSF solutions during dopamine application experiments.

### General two-photon imaging methods

Two-photon imaging was performed using a two-photon resonant-galvo scanning microscope (NeuroLabWare) at 15.5 frames/second and 796 x 512 pixels/frame as described previously (Lutas et al., 2019). An InSight X3 laser (Spectra-Physics) was used to excite the fluorophores (910-1050 nm), and the emission light was filtered (green: 510/84 nm; red: 607/70 nm; Semrock) before collection with photomultiplier tubes (H10770B-40; Hamamatsu). The XY scanning was performed using resonant/galvo mirrors and the Z scanning was achieved with an electrically-tunable lens (Optotune).

### Two-photon imaging of acute brain slices

For two-photon imaging of acute brain slices, slices were prepared as described in Lutas *et al.* and transferred to a recording chamber perfused with ACSF (oxygenated with 95% O_2_ and 5% CO_2_; flow rate: 2-5 mL/min) at either room temperature or 32° C as indicated. Imaging was performed with a 16x 0.8 NA water-immersion objective (Nikon). The excitation wavelength used was 920 nm.

In optogenetic experiments involving Chrimson, stimulation was triggered by the onset of a frames (15.5 Hz) for 2-, 5-, or 10-second duration. The gating property of the PMT was triggered at the onset of the frame to protect the PMT from optogenetic stimulation light and lasted for 10 ms. Thus, the top ~16% of each frame during the stimulation was blank, but much of the frame (~84%) provided near-simultaneous imaging of biosensor signals during the photostimulation. A 620 nm LED (1 mW/mm^2^, Luxeon Star LEDs) driven by an Arduino-controlled driver (Luxeon Star LEDs) was used for photostimulation.

In optogenetic experiments involving biPAC, each slice was imaged for 10.5 minutes (15.5 frames per second). At time points: 0.5 min, 2.5 min, 4.5 min, 6.5 min the PMT was powered off, a 470 nm LED (1 mW/mm^2^, Luxeon Star LEDs) driven by an Arduino-controlled driver (Luxeon Star LEDs) was then turned on for 2 seconds, and the PMT power was the restored to the original level. The PMT was turned off and on to protect it from the LED light.

To block phosphodiesterase activity, IBMX (100 μM in DMSO) was applied to the brain slice for at least 10 minutes. Recordings of cADDis following biPAC stimulation were performed before and after application of IBMX. To block dLight1.1 signals (dLight1.1 is engineered from the D1 receptor), we applied an antagonist of the D1 receptor (SCH23390; 300 nM) while recording dLight1.1 signals evoked by photostimulation of dopamine release from VTA^DA→BA^ axons in brain slices.

### *In vivo* two-photon imaging via implanted lens

Two-photon imaging via implanted lenses was performed as previously described (Lutas et al., 2019) with the following adjustments. A 10x 0.5 NA air objective was used (ThorLabs). For optogenetic stimulation of Chrimson via the implanted lens, a 617 nm LED (ThorLabs) was used (5 - 10 mW at the objective face).

### Fiber photometry recording

Fiber photometry recordings were performed as described in Lutas *et al.*, 2019, using head-fixed mice that were free to run on a circular treadmill. Fiber optic cables (1 m long; 400 μm core; 0.48 NA; Doric Lenses) were coupled to implanted optic fibers with zirconia sleeves (Precision Fiber Products). Excitation and emission light was passed through a four-port fluorescence mini-cube (FMC4_E(460-490)_F(500-550)_O(580-650)_S, Doric Lenses), which allowed for collection of GFP fluorescence and excitation of red-shifted channelrhodopsins. For biosensor photometry recordings, the excitation light (~ 100 μW at the face of the patch cord) was provided by a 465 nm LED (Plexon LED and driver). For optogenetic stimulation, the excitation light (~ 5 mW at the face of the patch cord) was provided by a 620 nm LED (Plexon LED and driver) which was controlled by an Arduino Uno. Emission light was collected by a femtowatt photoreceiver (Newport 2151), demodulated using a lock-in amplifier (SR830; Stanford Instruments) and digitized at 1 kHz sample rate (PCIe-6321; National Instruments). Data acquisition was controlled using a custom script in MATLAB (MathWorks). In a subset of experiments the D1 receptor agonist (SKF81297; 20 mg/kg in saline; Tocris R&D systems), was injected intraperitoneally while recording cADDis photometry signals. In these experiments, endogenous dopamine release was achieved with Chrimson photostimulation of VTA^DA→BA^ terminals and we compared peak cAMP responses following photostimulation and injection of SKF81297. Delivery of appetitive (Ensure) or aversive (tail shock; 0.3 mA,50 ms) outcomes was performed as previously described (Lutas et al., 2019).

## QUANTIFICATION AND STATISTICAL ANALYSIS

### Statistics

The numbers of samples in each group were based on those in previously published studies. Experiments were conducted by an investigator with knowledge of the animal genotype and treatment. Custom-written MATLAB analysis scripts allowed for data analysis in an automated and unbiased manner. All virus expression, optic fiber implants, and GRIN lens placements were verified by *post hoc* histology. Mice in which either the virus expression or optic fiber was not appropriately located (< 10 % of the time) were excluded from analysis. All data presented as bar and line graphs indicate mean ± s.e.m. with individual data points also plotted. Statistical analyses were performed in MATLAB. Significance levels are indicated as follows unless otherwise specified: *p < 0.05; **p < 0.01; ***p < 0.001.

### Data analysis

All data analyses were performed using MATLAB (Mathworks) and ImageJ (NIH).

### Fiber photometry analysis

Photometry signals were sampled at 1 kHz, low pass filtered below 100 Hz, and downsampled to 50 Hz. We calculated ΔF/F = (F − F0)/F0, where F0 the mean of baseline window prior to the event of interest. For analysis of the responses to an individual event, all trials containing presentations of that event during a run were averaged to obtain a mean timecourse, and then the peak response during the event window was obtained. We also did not use the sliding-window method for baseline normalized to avoid distorting the slow dynamics of intracellular cAMP signals.

### Widefield fluorescence brain slice imaging analysis

Movies were initially corrected for x-y motion using identical methods (efficient subpixel registration to averaged reference image) as used for two-photon imaging analyses above. For cADDis cAMP sensor imaging, as cells were very bright at baseline (as evoked cAMP leads to a decrease in fluorescence of this sensor) and showed slower dynamic changes in fluorescence upon dopamine application, regions-of-interests were automatically segmented by using morphological filters to identify bright spherical regions (Liang et al., 2018). Briefly, a mean projection through the movie was first applied to obtain a single mean image. Local normalization was then applied by subtracting the local mean (Gaussian kernel with sigma = 8 pixels) and dividing by local variance across pixels (Gaussian kernel with sigma = 150 pixels). Basic MATLAB functions were then used to remove small unconnected structures and fill in gaps in larger structures. The image was then binarized and regions of interest were segmented by applying a Euclidian distance transform followed by a watershed transform (a common strategy used to segment spherical objects). We estimated neuropil signals by taking circular annuli around the region of interest, as described above for two-photon image analysis.

### Two-photon brain slice and *in vivo* imaging analysis

Image registration for brain slice and *in vivo* two-photon calcium imaging of BA cell bodies was performed as previously described (Lutas et al., 2019). Briefly, to corrected for x-y motion, each frame from an imaging session was registered to a reference image (average of 1000 frames within a session) using efficient subpixel registration methods (Bonin et al., 2011). For extraction of signal from cell body regions of interest (ROIs) from volumetric brain slice imaging (15 depths; ~ 10 μm apart), we used CellPose (Stringer et al., 2020), which optimally identified ROIs from bright cADDis expressing cell bodies. For *in vivo* calcium imaging analysis, we used PCA/ICA to extract masks of pixels with correlated activity, corresponding to individual axons or cell bodies (Mukamel et al., 2009). Timecourses were extracted by averaging each of the pixels within each binarized mask. We calculated neuropil activity as the median value of an annulus surrounding each ROI (inner radius: 15 pixels; outer radius: 50 pixels; pixels belonging to any other ROI were excluded from these annulus masks). This timecourse of neuropil activity was then subtracted from the activity timecourse of the associated ROI to create a fluorescence timecourse, F(t), where t is time of each imaging frame. The change in fluorescence was calculated by subtracting a running estimate of baseline fluorescence (F0(t)) from F(t), then dividing by F0(t): ΔF/F(t) = (F(t) − F0(t))/ F0(t), where F0(t) is a running estimate of baseline fluorescence calculated as the 10^th^ percentile of F(t) in the previous 32-second sliding window (Petreanu et al., 2012).

### Criteria for determining responsivity to cues from *in vivo* calcium imaging

To determine if a cell was responsive to each cue, we used previously established, conservative criteria (Lutas et al., 2019), which are described here. We performed a Wilcoxon sign-rank test for each frame post-stimulus onset against the 1-s baseline period prior to stimulus onset, with Bonferroni correction for multiple comparisons across frames (p < 0.01). If three consecutive frames were significantly different than the baseline period, a cell was considered responsive to that cue. For all cells with significant responses to at least one cue, a cell preferred cue was determined as the cue evoking the largest response during the cue period. For estimation of a cell’s mean cue-evoked response magnitude, and for estimation of a cell’s response bias to a given cue, we averaged all trials containing presentations of that cue during the run to obtain a mean timecourse for that cell, and then the maximum response during the 2-s duration of the cue presentation was used as that cell’s response magnitude.

### Modeling dopamine-evoked cAMP dynamics

To model the dynamics of dopamine and cAMP, we first fit monoexponential functions to averaged traces of either dLight1.1 or cADDis recordings. We used the decay rate from the dLight transient to first model dopamine dynamics in response to 2 s square inputs (30 s intervals between each input), which exponentially depressed such that the second input was 25% weaker than the first pulse (as observed in our measurements of dopamine synaptic depression). We convolved this input kernel with an exponential (4 s time constant; step size of 0.1 s). We then used this convolved waveform which models dopamine with synaptic depression as an input kernal to model the cAMP dynamics (20 s time constant, step size of 0.1 s). For qualitative comparison with *in vivo* recordings, we normalized the modeled cAMP dynamics to the peak of the first evoked response.

